# Mechanistic insights into the association and activation of the SARS-CoV-2 2’-O-Methyltransferase (NSP16)

**DOI:** 10.64898/2026.04.15.718757

**Authors:** Heng Ma, Alexander Brace, Monica Rosas-Lemus, S. Chakra Chennubhotla, Karla J. F. Satchell, Arvind Ramanathan

## Abstract

The nsp16 2’-O-Methyltransferase is an essential non-structural protein of SARS-CoV-2, which methylates the viral mRNA cap structure, enabling it to evade the host immune response for higher translation efficiency. However, nsp16 is only active when it is bound to its cofactor, namely the non-structural protein-10 (nsp10). Understanding how nsp10 binds to and activates nsp16 function can help to develop targeted inhibitors; however, given the varying degree of disorder in both nsp10 and nsp16, characterizing this interaction has been challenging. Using long-timescale molecular dynamics simulations and AI/ML methods, we posit that the nsp16/nsp10 binding process is mediated by a hydrophobic latch formed with Leu4298 from nsp10 and a hydrophobic concave on the nsp16 protein surface. Our study highlights how the nsp16 S-adenosyl-L-methionine (SAM) pocket closes in its monomer state, which in turn deactivates the MTase function. We also observe that the nsp16/nsp10 complex allows for the RNA binding site to open with the empty SAM pocket. The results reveal how the SAM pocket loops facilitate SAM binding while allowing for the by-product S-adenosyl-L-homocysteine (SAH) to exit. Our study thus provides valuable atomistic-level mechanistic insights into understanding the activation of nsp16 MTase function while highlighting the challenges of studying protein-protein interactions mediated by largely flexible/disordered regions.

**SIGNIFICANCE:** Nsp16 carries out the methylation of the viral mRNA to gain immune evasion and translation efficiency. Understanding its complex molecular machinery can help us develop better therapeutic treatments. Here, we explore the key activation conditions for the SARS-CoV-2 nsp16 function via molecular dynamics simulation and AI/ML methods. The results demonstrate the role of nsp16 loops in different stages preparing for the methylation reaction from nsp16/nsp10 binding, (de)activation of nsp16 function and how the nsp16 SAM binding pocket can affect the RNA binding loops. This research explains the role of the nsp16 loops, which orchestrate its molecular function, and provides valuable insight to develop more targeted therapeutic approaches to disrupt viral immune evasion activity.

## 1 INTRODUCTION

The past coronavirus pandemic (COVID-19) resulted in over 7 million deaths and 676 million infections, resulting in tremendous human, social, and economic costs, as of 03/10/2023 – the last time when these reports were made publicly available (1). Though the number of infections and hospitalizations due to COVID-19 has dropped due to the availability of effective vaccines, it surged in many states as of 08/2024 (2). To better prepare for new variants and other novel highly pathogenic coronaviruses, it is important to understand the conserved mechanisms of viral replication. This understanding can provide strategies for therapeutic interventions to target future zoonotic coronavirus outbreaks (3, 4).

Of the various SARS-CoV-2 viral proteins, several non-structural proteins (nsps) play important roles in the viral life cycle, offering potentially novel ways to develop therapeutics targeting them (5–7). Of these nsps, the 2’-O-methyltransferase (MTase), referred to as nsp16, plays a vital role in evading the host immune response by mimicking the function of the Cap-specific mRNA (nucleoside-2’-O-)-methyltransferase (CMTr1) (8, 9). Nsp16 is an enzyme that transfers a methyl group from the cofactor, S-adenosylmethionine (SAM) to the 2’-hydroxyl group of the ribose sugar of the first nucleotide of the viral mRNA cap structure, converting the Cap-0 structure (m^7^GpppN) to a Cap-1 structure (m^7^GpppNm). This methylation allows better translation efficiency of the viral mRNA and enables the virus to evade the immune response by not letting pathogen recognition receptors bind to the viral mRNA (10–12). Experiments have demonstrated that knocking out nsp16 function can inhibit viral growth, thus offering a potential strategy to target this enzyme with small molecules (13–15).

One of the unique aspects of the SARS-CoV-2 nsp16 is that it forms a functional heterodimer with nsp10 (Figure 1) (10, 11, 16–20). The nsp10 modulatory subunit is essential for the SARS-CoV-2 nsp16 to engage its MTase activity. Interestingly, nsp16 cannot bind to its substrate SAM (or RNA/ other ligands) without nsp10, indicating nsp10’s role as a stabilizer of the highly dynamic nsp16 binding site (11, 12, 18). The nsp16 structure consists of a classic Rossmann, *β*–sheet fold sandwiched between 11 *α*-helices and 20 flexible loops. These loops construct the binding pockets for both mRNA and SAM cofactor, as well as the binding interface with nsp10. Among them, four particular loops, namely, the gate loops GL1 (residue 6817-6840, shown in red in Figure 1) and GL2 (residue 6928-6942 green in Figure 1) are involved in stabilizing the RNA substrate within the active site, while SAM-binding loops SAMBL1 (residue 6869-6875, dark blue in Figure 1; also known as the coronavirus insertion loop (21)) and SAMBL2 (residue 6897-6901, magenta in Figure 1; metal binding loop (21)) are involved in the SAM cofactor binding pocket (17, 18). Notably, SAM forms strong hydrophobic interactions through the adenosine group within the pocket, followed by hydrogen bonds with the binding loops. The GL loops can also adopt different poses, accommodating the RNA binding process (17).

**Figure 1:**
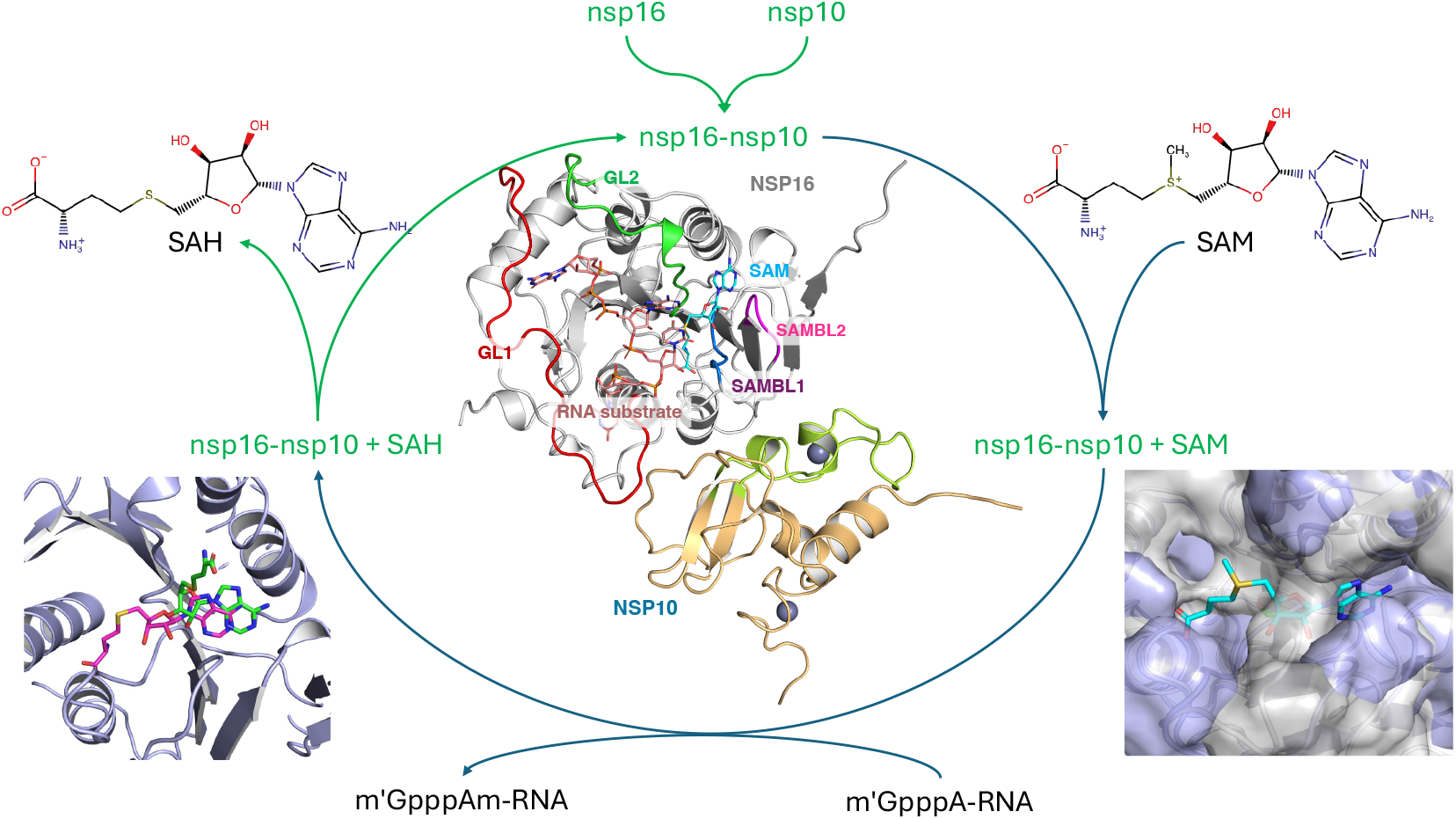
The overview of nsp16/nsp10 protein complex MTase function. The simulated processes are colored in green. (center) The overview of nsp16/nsp10 heterodimer complex with nsp16 colored in silver, nsp10 colored in light orange, and SAM molecule (cyan). The binding loops of nsp16 are highlighted as SAM binding loops, SAMBL1 in dark blue and SAMBL2 in magenta, and the mRNA binding loops, GL1 in red and GL2 in green. The m^7^GpppA mRNA (salmon) is from 7JYY PDB entry and overlapped to the RNA pocket. The peptidomimetics region of nsp10 is colored in lime. The SAM binding pocket is sampled binding with SAM (bottom right) or SAH (bottom left). The SAH exited the pocket during the simulation, as the product of the MTase process.

The modulatory subunit, nsp10 is a 148-residue protein composed of two zinc fingers and interacts with both nsp16 and nsp14 (3’-5’ exoribonuclease) and is part of the larger replication-transcription complex (RTC) (16, 22, 23). Nuclear magnetic resonance studies have shown nsp10 to be a highly dynamic protein, able to undergo significant conformational changes (24). Its structure was resolved in monomer form and multimer, binding with nsp16 and nsp14 (17, 23, 25). It comprises a long unstructured loop (residue 4288-4317) around the center antiparallel, *β*-sheets. Upon binding with nsp14 or nsp16, the loop rearranges itself and sits along the binding interface with nsp16 (17, 23, 25). The protein contains two zinc cations bound with histidine and cysteine clusters, which are essential to stabilize the local protein structure and possibly to stabilize the bound m^7^GpppNm RNA substrate. (17)

Recent studies, including multiple sequence alignment, X-ray crystallography of mutant nsp10 structures and metadynamics simulations have shown that nsp10 is critical for stabilizing nsp16’s active conformation (26). Further, the authors posit that the nsp10 binding facilitates MTase activity by enabling proper SAM binding in nsp16, but that nsp10 mutants don’t necessarily affect this process (26). Further, nsp16/nsp10 consists of multiple binding sites for small molecules, as demonstrated recently where adenosine analogs demonstrated MTase activity inhibition (27). However, the dynamics, function and potential targetability of these binding sites remain unknown. Despite the presence of cryptic binding pocket (18), and other druggable sites, designing potent MTase inhibitors has remained challenging. We posit that a detailed mapping of conformational states involved in nsp16/nsp10 binding process could provide much needed insights in targeting nsp16.

Therefore, we set up molecular dynamics (MD) simulations to model nsp16/nsp10 binding process, while taking into account the presence of various small molecules, including the methyl donor SAM, reaction product S-adenosylhomocysteine (SAH), and MTase inhibitor sinefungin (SFG) (Table 1). We then systematically compare the conformational changes that are unique to nsp16 monomer and the nsp16/nsp10 heterodimer, especially on the nsp16 SAM pocket. We next compare the effect of three ligands on nsp16 binding loops, namely SAM, SAH and SFG, with the apo-nsp16/nsp10 system. These simulations highlight a distinct behavior of the GL1 and GL2 loops, affected by the state of SAM pocket. We use novel machine learning techniques to characterize a series of intermediate state ensembles that allow the two proteins to bind. We posit that the interface of nsp16/nsp10 is largely mediated by a single hydrophobic latch (Leu2498 of nsp10) scaffolded by unstructured loops within nsp16. Taken together, our study provides insights into the coupled dynamics of the network of binding sites within nsp16/nsp10, which suggests novel directions for targeted therapeutic design.

**Table 1:** Molecular dynamics simulation setups.

| System label | Ligand | PDB code | Simulation Setup ($\mu s$) | Total ($\mu s$) |
| --- | --- | --- | --- | --- |
| **Heterodimer simulations** |  |  |  |  |
| *comp* | None | 6W4H<sup>a</sup> | $0.5 \times 3$<sup>b</sup> | 1.5 |
| *comp_sam* | S-adenosylmethionine (SAM) | 6W4H | $0.5 \times 3$ | 1.5 |
| *comp_sah* | S-adenosyl-L-homocysteine (SAH) | 6WJT | $0.5 \times 3$ | 1.5 |
| *comp_sfg* | Sinefungin (SFG) | 6WKQ | $0.5 \times 3$ | 1.5 |
| **Monomer simulations<sup>c</sup>** |  |  |  |  |
| *nsp10* | N/A | 6W4H | $0.5 \times 3$ | 1.5 |
| *nsp16* | no-ligand, SAM, SAH, SFG | 6W4H | $0.5 \times 3 \times 4$<sup>d</sup> | 6.0 |
| **Binding simulations** |  |  |  |  |
| *comp* | None | 6W4H | $0.2 \times 20 \times 3$<sup>e</sup> | 12 |

<sup>a</sup> SAM is removed from the crystal structure.
<sup>b</sup> The simulation length is labeled as $L_{sim} \times N_{rep}$ , where $L_{sim}$ is the length of a single simulation and $N_{rep}$ is the number of replicas. For example, $0.5 \times 3$ means 3 runs of 500-ns MD simulations. For the *comp* systems, we also ran 20 200-ns simulations of nsp16/nsp10 separated at various distances.
<sup>c</sup> The monomers are set up with the crystal structure of the heterodimer.
<sup>d</sup> The *nsp16* is run with 4 ligand-binding states, and each is with 3 replicas.
<sup>e</sup> 20 runs of simulations with nsp16/nsp10 separated at different distances, and each is run with 3 replicas.

**Table 2:** The binding free energy of nsp16/nsp10 complex in different binding states from MM/PBSA calculation in kcal/mol.

| State | $R_{com}$ (Å) | $N_{ca}$ | $E_{vdw}$ | $E_{EL}$ | $E_{PB}$ | $E_{NP}$ | $\Delta G_{gas}$ | $\Delta G_{solv}$ | $\Delta G_{total}$ |
| --- | --- | --- | --- | --- | --- | --- | --- | --- | --- |
| Native Binding | 32.0 | 151.3 | -93.2540 | -212.8038 | 267.8414 | -9.6352 | -306.0578 | 258.2062 | -47.8516 |
| Transition-1 | 34.2 | 78.8 | -41.4122 | -204.5802 | 220.7516 | -5.0959 | -245.9924 | 215.6557 | -30.3367 |
| Transition-2 | 38.1 | 33.5 | -7.6494 | -128.4983 | 129.6347 | -1.1529 | -136.1477 | 128.4819 | -7.6659 |
| Separate | 56.8 | 4.0 | -0.1477 | -48.2772 | 47.8281 | -0.0093 | -48.4249 | 47.8188 | -0.6061 |

## METHODS

We used MD simulation as the primary tool for sampling the molecular conformations. Root-mean-square fluctuation (RMSF) of the protein backbone was observed to identify the domain of interest during the MD simulation. The trajectories from MD simulations were also compressed into low dimension latent representation using the Autoencoder machine learning (ML) framework and subsequently clustered into distinct conformational states with MinibatchKmeans and modularity optimization. We also calculated the binding free energy of nsp10 and nsp16 using Molecular Mechanics/Poisson-Boltzmann Surface Area (MM/PBSA) in different binding states. Moreover, the MD trajectories captured molecular conformations of nsp16 under different binding conditions with nsp10 and various ligands, which provided an atomsitic insights of how the unstructured loops modulated and performed in its enzymatic function.

### Molecular dynamics simulations

MD simulations were set up to model nsp16 in various binding states with nsp10 and SAM-like small molecules. In total, we ran 25.5-*µs* of simulations on Nvidia V100 GPUs (Table 1).

#### Probing the equilibrium conformational fluctuations of the nsp16/nsp10 complex

The simulations were carried out using the OpenMM high performance simulation toolkit (28) on CUDA platform with mixed precision. Molecular models were built using the crystal structures from PDB data bank, as shown in Table 1 (17). We built a total of four systems of the nsp16/nsp10 complex, based on the ligand in the binding site, *comp* as apo-nsp16/nsp10, *comp_sam* as nsp16-SAM/nsp10, *comp_sah* as nsp16-sah/nsp10 and *comp_sfg* as nsp16-SFG/nsp10. Both initial structures of *nsp16* and *nsp10* monomers were taken from the 6W4H PDB entry directly. Therefore, the nsp16 run was simulating the deactivation process instead of the activation upon binding.

For each of the systems, atomic interactions were described with Amberff99sb-ildn force field (29). After determining the protonation states of each of the residues at pH 7.0, appropriate counter ions were added to neutralize the systems (nsp16 and nsp10 have a charge of −7 and +4 respectively). The ligand parameters were generated using the antechamber program with General AMBER Force Field (GAFF) (30, 31). The binding cysteine and histidine residues to the zinc-finger architecture of nsp10 were deprotonated to retain proper binding with the cations. The systems were solvated with explicit TIP3P water models using a salt concentration of 0.15 M NaCl. Nonbonded interactions were cut off at 10 Å with periodic boundary condition (PBC). Particle Mesh Ewald method (32) was used to calculate long-range Coulombic interactions. The simulation was integrated with Langevin integrator with 2 fs timestep and 1 ps^−1^ heat bath friction coefficient in 310 K. The system pressure was maintained at 1 bar with a Monte Carlo barostat. Each MD run reported data every 50 ps. Root mean square fluctuation was calculated from the MD trajectories and used to identify the interesting regions that drove molecular state transition.

#### Probing the binding conformational landscape of nsp10 and nsp16

To characterize how nsp10 and nsp16 interact with each other, we simulated the complex formation process by allowing the two proteins to approach each other. Within the crystal structures of the nsp16/nsp10 complex, the distance between the centers of mass (CoMs) of nsp10 and nsp16 was ∼ 32 Å. Conformers of unbound nsp10 and nsp16 were obtained by pulling them away from each other. We generated a total of 20 configurations up to 10 Å separation with 0.5 Å increment. Each of these structures was then equilibrated (as described above) and then simulated under the same conditions for 200 ns with 3 replicas.

### Molecular Mechanics/Poisson-Boltzmann Surface Area (MM/PBSA) for binding free energy

The binding free energy of nsp16/nsp10 complex was calculated using MM/PBSA.py implementation from Amber (33). The free energy was separated into dry and solvent contributions. The dry contribution of the binding energy was directly estimated through molecular interaction terms from the MD simulation setup. The solvent contribution consisted of two terms, the polar term and nonpolar term. The former was obtained by solving the Poisson-Boltzmann equation in solution and vacuum environment with dielectric constant at 80 and 1, respectively. The ionic strength was set at 150 mM. The nonpolar term was affected by the solvent-accessible surface area (SASA) calculation, probed with 1.4 Å sphere and surface tension of 0.005 kcal/mol Å^2^.

### Machine learning (ML)-based characterization of the nsp16/nsp10 binding conformational landscape

#### Convolutional variational autoencoder (CVAE)

To characterize the binding conformational landscape of nsp16/nsp10 complex, we used an unsupervised machine learning (ML) approach that learns a latent representation of the conformational space. The conformers from nsp16/nsp10 complex were embedded into a low-dimensional latent space via a convolutional variational autoencoder (CVAE) (34) built with Keras/Tensorflow framework (35). Each nsp16/nsp10 dimer conformation was represented with C_*α*_ contact map between nsp10 and nsp16 with 10 Å cutoff distance. The CVAE was trained in an unsupervised fashion, by encoding the protein contact maps into a latent space and reconstructing the full contact maps from the low-dimensional embeddings. The convolutional layers allow the autoencoder to focus on local patterns in the image, and the variational layer at the bottleneck regularizes the embeddings to conform to a normal distribution in the latent space. Using contact information, the CVAE can embed dimer binding poses in the latent space, based on the conformational similarity. The 252,000 input contact maps are shuffled and split with 80/20 training/validation ratio. Ten CVAE networks were built with 7 different latent dimensions (3, 5, 10, 20, 30, 40, 50, and 70) to identify the best latent representation, and trained on Nvidia V100 GPUs. A latent dimension of 50 was selected to represent the 3D complex conformations by considering the lowest validation loss across the training experiments.

#### Cluster and Modularity Analysis

Furthermore, we employ clustering methods to categorize the conformers from MD simulations into different conformational states of the protein binding process, taking advantage of the low-dimensional CVAE embeddings. We utilize the MiniBatch Kmeans clustering method from scikit-learn to organize the CVAE embeddings into 200 clusters (36, 37). The clusters were then treated as nodes in a graph, connected with edges representing the state transitions observed in the MD simulations. Ideally, in this network each node represents a distinct molecular substate that is connected to neighboring substates according to the presence of a state transition. However, the K-means method arbitrarily divides the latent embeddings into the designated number of states, which needs to be optimized or otherwise introduces bias to the clustering results. To eliminate this artifact of K-means, we carried out a modularity optimization on the graph, which is designed for this kind of classification task (38, 39). It compresses the original graph by iteratively combining highly interconnected clusters until point modularity is maximized. From a biophysical perspective, the molecular states that have a high transition rate between them are merged into a single state. As a result, we obtain a smaller 41-node graph which meaningfully captures the physical information from the MD simulations.

### Nsp16 binding pocket analysis

#### Loop backbone dihedral

To understand how nsp16 carries out its MTase function, we characterized and measured the occlusion space of the SAM binding pocket. Both experimental and simulation results from previous studies indicate that the presence of nsp10 is essential for SAM pocket to remain open (11, 12, 18). We tracked the backbone dihedral angles of the binding loops to understand the change between nsp16 monomer and heterodimer states.

#### SAM pocket volume base analysis

Residue-based characterization is however insufficient in capturing a complex mechanism, where multiple residues are involved. Therefore, we measure the size of the binding pocket via a grid space analysis to directly quantify the effect of nsp10 binding to the SAM binding pocket of apo-nsp16. For each conformer, an imaginary SAM molecule is placed into the binding pocket, by aligning it to the 6W4H crystal structure. The occlusion space is defined as the overlapping volume between the nsp16 protein and this SAM molecule, and calculated by slicing the molecular space into 0.25 Å cubic grids. For each atom, its radius is calculated using the force field Lennard Jones parameter, *σ*, and its occupied space is defined by grids, whose centers are within the radius to its center. The grids that were in both protein and ligand volume are the occlusion space. For example, the 6W4H structure has a small overlapping of 3.91 Å^3^ in volume between nsp16 and SAM.

## RESULTS

### SAM binding pockets of nsp16 closes in monomer state

Both experimental and simulation results have shown the SARS-CoV-2 nsp16 is inactive without the nsp10 co-factor. (12, 16, 18) However, there is a lack of atomistic level understanding of why nsp16 is not active in its monomeric state. To investigate this question, we compared the apo-nsp16 protein in monomer form and in the heterodimer form bound with nsp10.

We first compared the nsp16 RMSF between the monomer and complex form, as shown in Figure 2. Clear differences were observed, where the monomer form demonstrated higher fluctuating loops around the binding interface region, underlined in purple in Figure 2A. As the nsp16 binding loops were overlapping or in direct vicinity of the interfacial ones, the differences also impacted the behavior of the nsp16 binding pockets, especially the SAM pocket. The GL2 was further away from the interface, but it constructed the opposite end of the SAM pocket to the SAMBLs, thus was affected by the binding indirectly as well. From the RMSF, we learned the nsp10 binding modulated the binding interface and led to more stable binding pockets. Without nsp10, the nsp16 demonstrated more flexible structures around the binding sites. However, we were still missing how these effects manifested the deactivation of nsp16 function. To directly quantify the effect of the flexible loops in the nsp16 monomer, we hypothesized the destabilized loops can no longer maintained a suitable binding pocket accommodating the SAM molecule, and designed a method to measure the occlusion between the protein and the SAM binding site. The method recorded the overlapping volume between the nsp16 and the space that a binding SAM molecule would occupy, given a nsp16 conformer.

**Figure 2:**
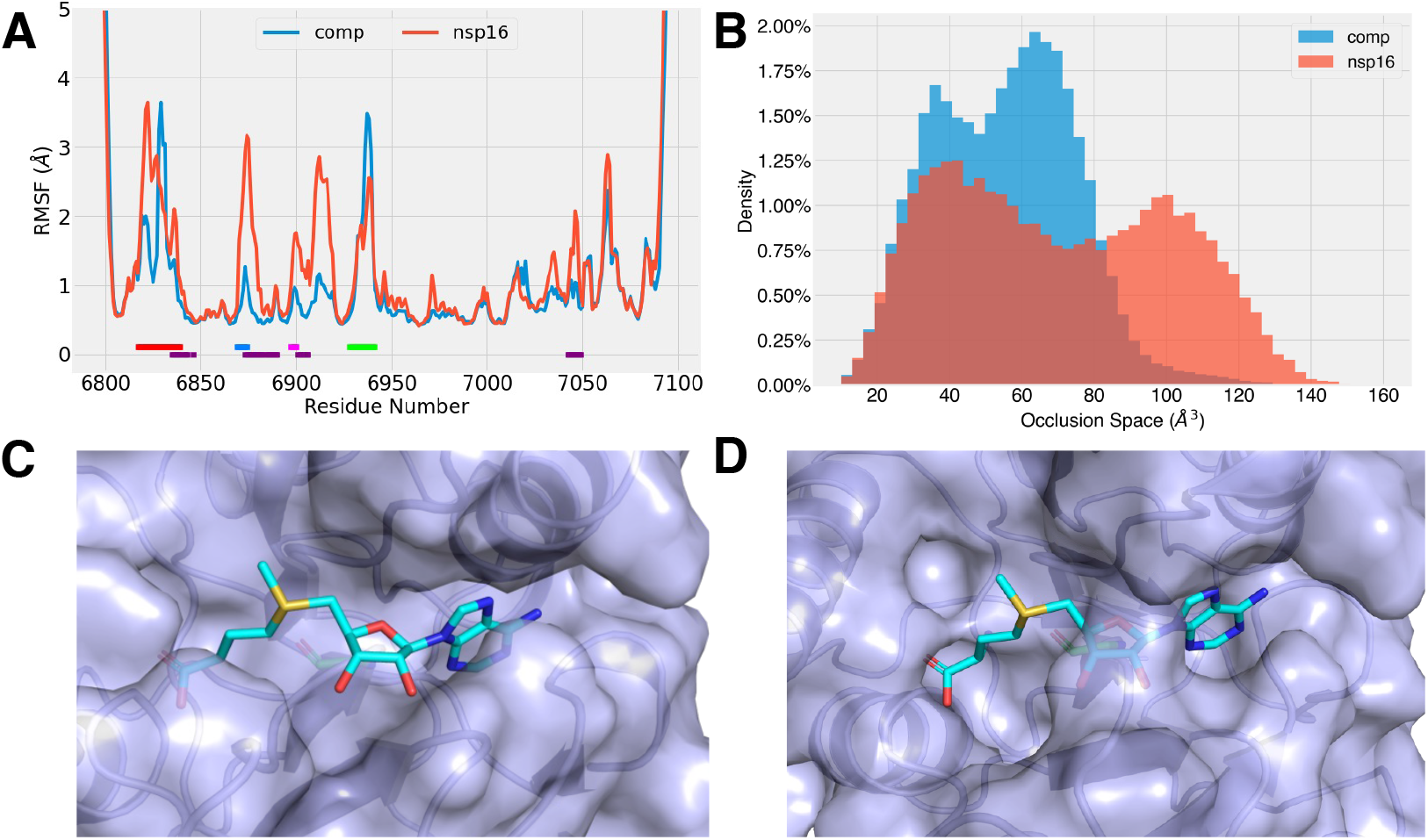
(A) The RMSF of nsp16 under monomer and nsp10-binding complex states. The nsp16 binding loops are underlined, as SAMBL1 in blue and SAMBL2 in magenta, and the mRNA binding loops, GL1 in red and GL2 in green. And the contacting regions are also underlined in purple. (B) Occlusion space between protein and SAM binding pose in nsp16 SAM binding sites under monomer and heterodimer conditions. (C-D) The open (C) and closed (D) SAM pocket in nsp16/nsp10 heterodimer and nsp16 monomer forms, respectively. The protein is shown as light blue surface and cartoon, and the SAM molecule in cyan is placed in the pocket for demonstration purpose. The open state (C) was observed with less overlapping between protein (blue surface) and SAM molecule (cyan sticks). In the closed state (D), the SAM molecule binding space was occupied by the protein.

The histograms of the occlusion space were shown in Figure 2B for both the monomer and nsp16/nsp10 dimer. Notably, the *6W4H* structure has occlusion space of 3.5 Å^3^. The complex with filled SAM pocket was with around 20 Å^3^ occlusion volume, shown in Figure S2. Both the nsp16 monomer and heterodimer simulations were initiated from *6W4H* (Table 1), where the SAM pockets were open in the initial state (Figure 2C). In both setups, the histograms of the occlusion space were in two-peak bimodal form. The low-occlusion state was observed at a similar range for both systems, ∼ 37 Å^3^ for the dimer system and ∼ 40 Å^3^ for the monomer. To give a perspective, the volume of a water molecule is around 29.7 Å^3^ (40), and that of a SAM molecule is 326.6 Å^3^. For the higher occlusion states, the nsp16 monomer was at ∼100 Å^3^ occlusion between the protein and SAM binding pose, and a lower occlusion of ∼65 Å^3^ was observed while binding with nsp10.

From our experiments, regardless of monomer or dimer states, the empty nsp16 SAM pockets shrank into smaller sizes, shown as in Figure 2B. Without the nsp10 modulation, the SAM pocket collapsed into a further closed state. The observation not only confirms, but also provides some quantified insights for previous findings that the nsp16 MTase function is disabled without the nsp10 modulation. (12, 16, 18) By examining the MD trajectories, we also identified the conformers, in which the SAM binding site is closed, as shown in Figure 2D. The binding pocket collapsed with the binding loops and moved into the binding space.

We further explored the mechanism of pocket closing in the nsp16 monomer form. The backbone dihedral angles, *ϕ* and *ψ* of SAMBL residues, were measured for the simulation runs of nsp16/nsp10 complex and nsp16 monomer. No difference was observed in SAMBL2 between the simulations of nsp16 monomer and complex form (Figure S3). The SAMBL1 showed more significant differences, and the results were shown in Figure 3A. SAMBL1 residues, 6868-6872, showed two different states in nsp16 monomer form, and the nsp16/nsp10 complex had a single state in the histogram. The additional state in the nsp16 monomer was identified with SAM pocket in closed form. By looking closer to those residues, the loops turned from the open state around the SAM pocket to the closed state, where the loop was overlapping with the SAM binding pose, shown in Figure 2D and Figure 3B. The residue Gly6869 was found to play a crucial role in the conformational change. Its *ϕ* angle shifted from 70° to 180°. Adapting to the change, the backbone around the SAMBL1 reached into the binding pocket and blocked SAM binding space.

**Figure 3:**
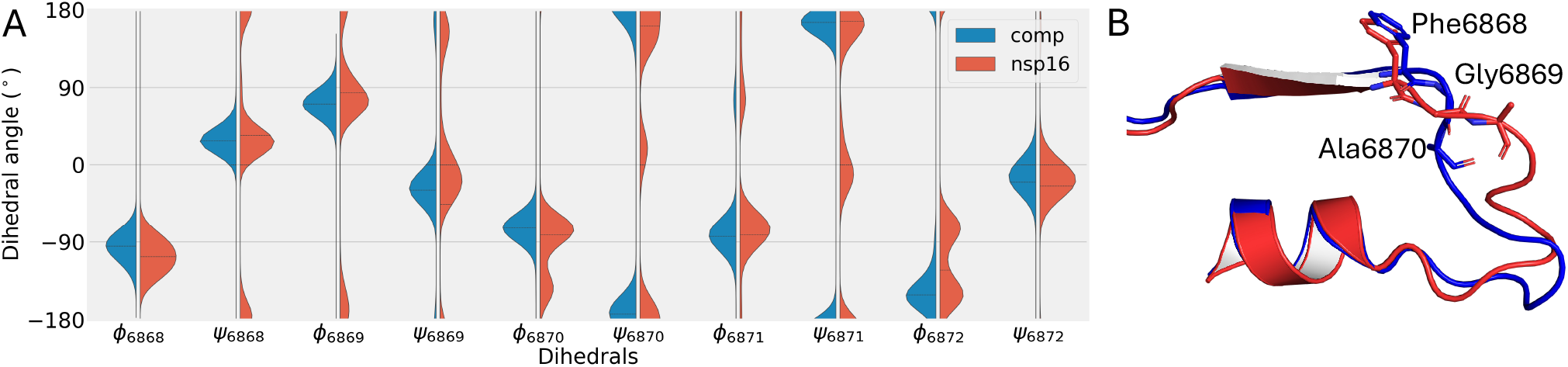
(A) The histogram plot of backbone dihedral angles of SAMBL1 residue 6868-6872, with the complex form in blue and nsp16 monomer in orange. (B) The 3D structure of SAMBL1 of nsp16/nsp10 heterodimer in open state (blue) and nsp16 monomer in closed state (red). The residue Gly6869 and its neighbors were labeled.

Additionally, we set up simulations of nsp16 monomer with binding SAM molecule as the initial conformation. During the simulation, the SAM molecule was observed to exit the binding pocket during the simulation. The SAM exiting process was recorded in the MD trajectory in the SI. The process was also confirmed with the occlusion histogram as well (Figure S2). This result confirmed that the nsp16 monomer lost its MTase function with an atom-scale mechanistic explanation.

### Empty nsp16 SAM pocket opened RNA site

In this section, we investigated how the binding ligand affected the nsp16/nsp10 complex conformation. Simulation runs were set up in the apo state, along with three different binding ligands (Table 1). The selected ligands represented different types of binding molecules during the MTase process, with SAM as the reactant, SAH as the product and SFG as the inhibitor. All three ligands were in a similar form, where the adenosyl group was connected to different amino acids (methionine for SAM, cysteine for SAH and L-ornithine for SFG).

The RMSF profiles of nsp16 in heterodimer complex form were calculated from the MD trajectories and shown in Figure 4A. The SAM pocket region showed similar fluctuation, even with apo-nsp16/nsp10 complex, for the region was already stabilized with nsp10 binding, as discussed previously. The main difference was observed at the RNA binding loops, GL1 and GL2. Around the GL1, the apo-nsp16/nsp10 complex had a different fluctuation profile from other systems with binding ligands. From the RMSF, GL1 was separated into three sections, GL1-I(6817-6825), GL1-II(6826-6834) and GL1-III(6835-6840). The GL1-I had significant lower chain fluctuation in the apo-nsp16/nsp10 than the ligand-binding systems, while similar RMSF profiles were observed at other two GL1 sections. The GL1-II had high fluctuation and GL1-III was with a low chain fluctuation for all systems. There existed a large gap between chain fluctuation between the GL1-I and GL1-II sections in the apo-nsp16/nsp10 system. To accommodate this gap in the dynamics with GL1-I, the GL1-II had a completely different chain conformation in the apo-nsp16/nsp10 simulations.

**Figure 4:**
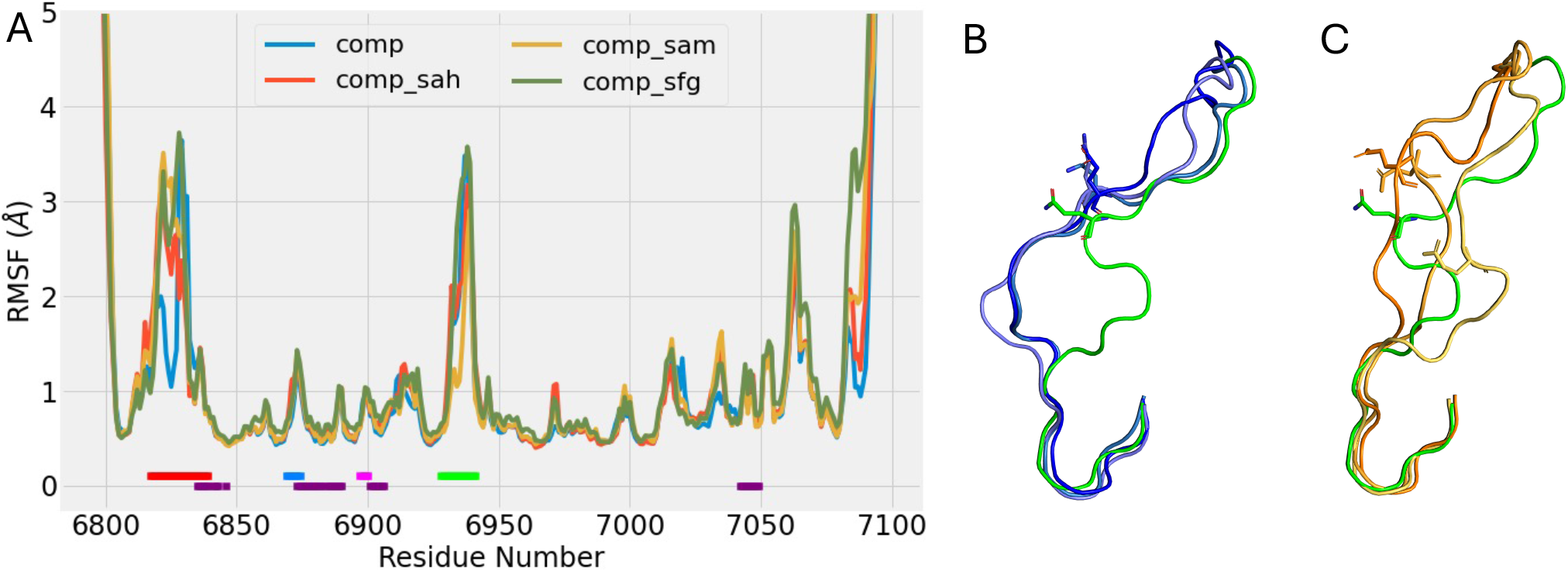
(A) The RMSF of nsp16 in nsp16/nsp10 heterodimer with different binding ligands in SAM pocket. The ligand binding and interfacial loops were underlined in the same fashion as Figure 2. (B) The opening of GL1 from the initial conformation (green) to the relaxed pose after simulation (blue) during 3 replicas of apo-nsp16/nsp10 simulation runs. (C) The conformations of GL1 loops after 3 replicas of nsp16-SAM/nsp10 simulation runs (colored in orange). The initial conformation is colored in green.

To further investigate the chain formation, we plotted the GL1 backbone dihedral angles of apo-nsp16/nsp10 and nsp16-SAM/nsp10. (Figure S4). The main difference was observed in the GL1-II section, where the backbone dihedral demonstrated distinctive behaviors, indicating the two systems were in different conformational states. All the apo-nsp16/nsp10 systems agreed on the conformation, where the GL-II section moved away from the RNA binding region, and availed the binding access to the region (Figure 4B). In the SAM-binding systems, the chain exhibited random conformation, amongst which no definitive chain configuration was observed from the simulation replicas (Figure 4C). Similar phenomena were found on other ligand binding systems (Figure S6). Therefore, the empty SAM pocket opened up the GL1 to facilitate the RNA binding.

The GL2 binds with both RNA and SAM in the solvated structure from crystallography. (17) It had similar dynamics with all systems, except for the nsp16-SAM/nsp10 simulations (Figure 4). The GL2 binding with SAM molecule showed a lower RMSF. While binding with SAM, the molecular motion of the GL2 were hinged around Thr6934 and Lys6935, which corroborates with the crystallography results (17). In the MD simulations, a hydrogen bond was observed between backbone N of Tyr6930 and O4’ of ribose from the ligands, stabilizing the neighboring regions. The donor-acceptor distances were observed around 3.1-4 Å. However, lower chain fluctuation was observed with SAM-binding system, as O4’ of SAM ribosome was found to be more negatively charged, from the partial charge estimation of AMBER semi-empirical quantum chemistry package, sqm (30, 41). As a result, the SAM binding stabilized GL2, comparing to SAH and SFG.

### SAH exits the binding pocket

The SAH molecule is the product of the MTase function, during which the methyl group transfers to the RNA. In our nsp16-SAH/nsp10 complex simulation, we observed the SAH molecule exited from the SAM pocket (Figure S7). The escaping of SAH from the pocket started with the amino acid portion moving out of the binding pose. We suspected the ligands had different flexibility, which resulted in different binding behavior. To confirm this hypothesis, we measured the bending angle of the amino acid and adenosyl groups as ∠(*C*)_*β*_ − *S*_*η*_ (*C*_*η*_) – *C’*, (Figure S8). By representing the distribution into a Gaussian mixture model, we found that all 3 ligands had a tri-modal distribution of this angle, but with different density. The SAH molecule had more even distribution among the 3 modes, while the SAM and SFG were confined into one or two modes. It indicated the SAH molecule was more flexible during the simulation, making it easier to escape from the SAM pocket in nsp16/nsp10 complex form. The inhibitor SFG emulated the mode of the reactant SAM and formed more stable interaction with the nsp16/nsp10 complex. The MTase enzyme utilized such feature of its reactant and product, to form a strong binding with the former, and allow prompt exit of the latter.

### Native binding state from MD simulation

Next, we investigated the forming of nsp16 and nsp10 complex. Starting with the native binding state of apo-nsp16/nsp10 heterodimer, the conformation from MD simulation runs is shown in Figure 5A. The structure of apo-nsp16/nsp10 remained stable with a RMSD of 2.05 Å to the *6W4H* crystal structure, and the secondary structures were also consistent with the crystallography results. (17) The heterodimer binding interface comprised primarily loops from both proteins. The nsp16 interfacial regions consisted of nsp16^*I*^ (residues 6835 to 6846), nsp16^*II*^ (6874 to 6889), nsp16^*III*^ (6900 to 6908), and nsp16^*IV*^ (7042 to 7046). These interfacial regions were also parts of nsp16 RNA/SAM binding mechanism. The nsp16^*I*^ region was part of RNA binding GL1 on the binding interface. The nsp16^*II*^ and nsp16^*III*^ are partially overlapping with nsp16 SAMBL1 and SAMBL2. For nsp10, we identified 3 interfacial regions at the interface, nsp10^*I*^ (4293 to 4300), nsp10^*II*^ (4322 to 4337), and nsp10^*III*^ (4346 to 4349). In the native configuration, nsp10^*I*^ loop fitted into the gap formed by nsp16^*I*^, nsp16^*II*^ and nsp16^*III*^. Nsp10^*II*^ and nsp10^*III*^ are part of peptidomimetics region, that binds with a Zn^2+^ cation.

**Figure 5:**
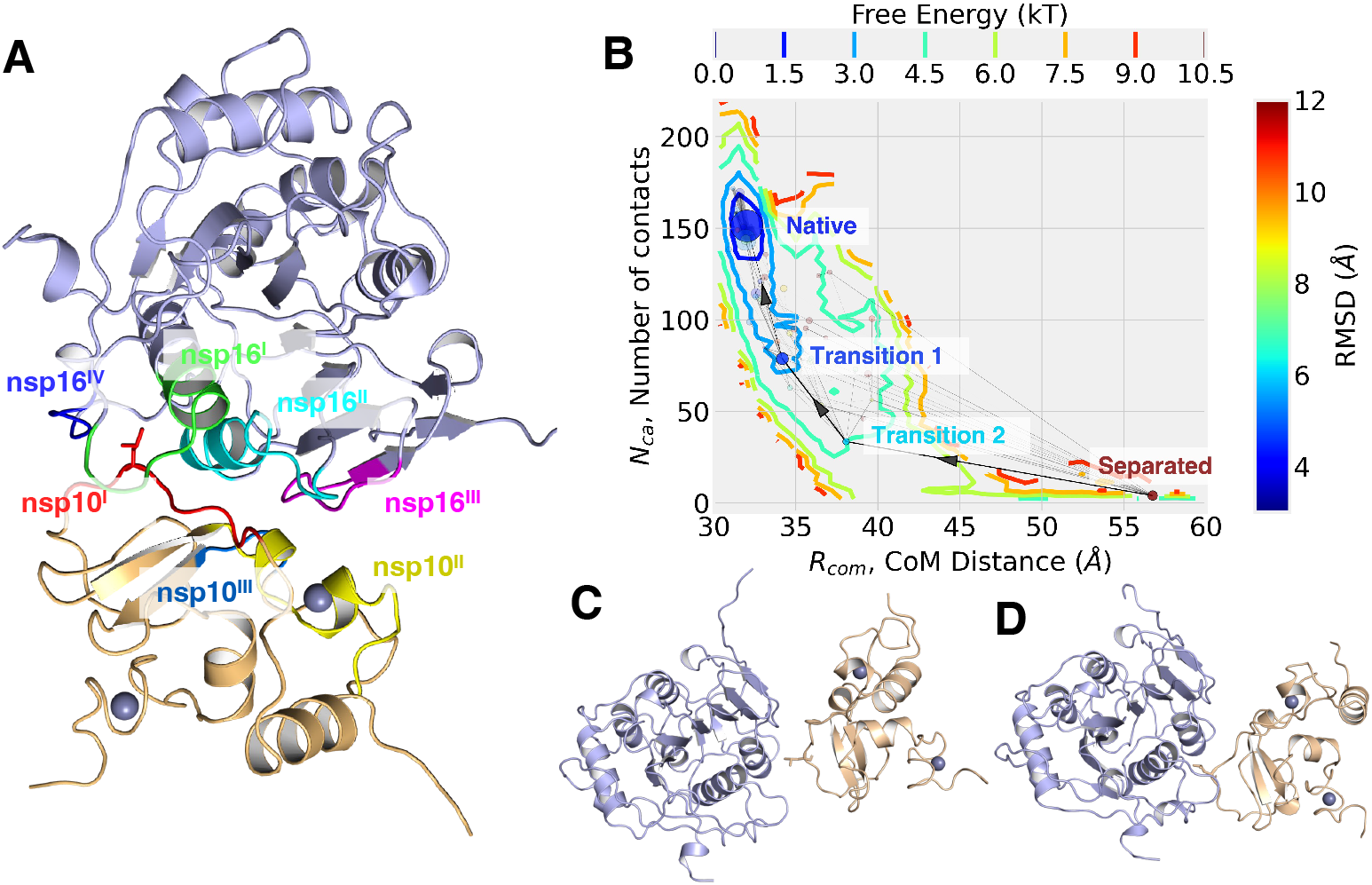
(A) The native states of the nsp16/nsp10 heterodimer (with nsp16 in light blue and nsp10 in wheat) from the MD simulation, with interfacial binding regions marked, nsp16^*I*^ in green, nsp16^*II*^ in cyan, nsp16^*III*^ in magenta, nsp16^*IV*^ in blue, nsp10^*I*^ in red, nsp10^*II*^ in yellow, nsp10^*III*^ in marine. The nsp10 Leu4298 on the binding interface is plotted in sticks. (B) The nsp16/nsp10 free energy interface according to the distance between nsp10 and nsp16 center of mass, and number of contacts between C^*a*^ within 10 Å. The configurational states are plotted with their average CoM distance and number of C_*a*_ contacts. Each one is colored based on its mean RMSD and its size based on the number of conformers in the state. The identified binding pathway is highlighted with solid lines and arrows. The separated, two transition and the native binding states are also marked. (C-D) The snapshot of transition state 1 (C) and 2 (D).

At the binding region, we identified a key residue, Leu4298 from nsp10^*I*^, which sit in a hydrophobic pocket, formed by nsp16^*I*^, nsp16^*II*^ and nsp16^*III*^ loops. The pocket consisted of hydrophobic residues, Pro6835, Ile6838, Val6842, Leu7042 and Met7045. Additionally, Leu4298 also formed a hydrogen bond between its backbone carboxyl O and Gln6885 side chain amide N_*η*_, at a distance of 2.7 Å. The Leu4298 played a crucial role during nsp16/nsp10 binding process, as discussed in the following section.

### A single hydrophobic latch facilitates formation of the nsp16/nsp10 complex

To explore the binding process, we picked out the conformers from the binding simulations, where the initial frames were with different separation between nsp16 and nsp10. The free energy surface was calculated with density of state, base on two selected observables, nsp16/nsp10 CoM separation distance, *R*_*com*_ and number of C_*α*_ contacts (10 Å cutoff), *N*_*ca*_, shown in **??**. The simulations sampled conformations *R*_*com*_ with between 29.8 and 60 Å, and in the range of 0-225. Hence, the energy surface covered both separated and contacted states. The native binding state, with the lowest binding free energy, was observed at *R*_*com*_ = 32.0 Å and *N*_*ca*_ = 155. Around the native binding regions (limited to ± 0.01 Å of *R*_*com*_ and ± 1 contacts of *N*_*ca*_), there are the 67 observed conformations from the MD conformers, and their average RMSD to the *6W4H* crystal structure was at ∼ 3.23 Å.

The free energy landscape revealed various binding states from the MD simulations, but it missed the dynamics of state transitions from the simulation. To further investigate the nsp16/nsp10 binding mechanism, we applied a series of dimension reduction and clustering methods to compress the MD trajectory conformers into different binding states and build a transition network connecting these states, base on the MD trajectories.

A CVAE is trained on the contact maps from MD conformers (34). We picked its latent space representations and clustered into 200 clusters with Mini Batch K-means (36, 37). We built a graph from these clusters by connecting them based on observed state transitions during MD simulation. These connections/edges were weighted on the number of observed transitions. However, the number of total states is arbitrarily assigned and led a bias towards the separation of conformational states. To remove the K-means bias, modularity optimization was applied to merge the highly connected nodes, indicating rapid state transition between the linked states. (38) From these analyses, we obtained 41 unique conformational states and connected them again with the edges that were weighted by the transition probabilities. The new smaller graph was then plotted in the free energy landscape on the average R_*com*_ and N_*ca*_ of each state/node in Figure 5B. This transition graph consisted of all the conformational states from our MD simulations. The nodes represented the separated molecular states, and the edges described the state transition between the linked nodes.

The graph clearly presented the nsp16/nsp10 binding pathway, and revealed the key states during the binding process (Figure 5). For the binding process, the separated state was with ∼ 56.9 Å CoM distance and ∼ 4 C_*a*_ contacts within 10 Å cutoff. The conformation is shown in Figure S1A. Additionally, along the binding pathway, two transition states were identified between the native binding and separated states (Figure 5C and D). The two proteins approached each other, and the average CoM distances reduced from the full separated ∼ 56.9 Å to ∼ 38.1 and ∼ 34.2 Å. More contacts were formed during the process, increased from 4 to 38 and 78.

The formation of the native binding state requires the specific orientation and interface. We noticed the nsp10 Leu4298 residue played a crucial role, modulating the binding process to secure the correct binding conformation. At the transition state 1, the nsp10 Leu4298 entered the nsp16 interfacial hydrophobic pocket at (Figure 5D). With the Leu4298 latched into the interfacial pocket, the interfacial loops rearranged to accommodate a more packed native binding state, where a hydrogen bond was formed between Leu4298 backbone carboxyl O and nsp16 Gln6885 side chain amide N_*η*_ to lock it in(Figure 5A). The complex pair adopted this unique latch-like formation to lock in the binding interface prior to the formation of native conformation.

We estimated the itemized binding energy between nsp10 and nsp16 in those four states using MM/PBSA. In their native states, the binding free energy between nsp16/nsp10 complex was −47.85 kcal/mol. It slowly decreased to ∼ 0 moving towards the separated state. From the itemized the energy contributions, we found the intermolecular interactions, van der Waals and electrostatic forces, as the main attraction term between two protein, and the solvent effects as the main energy barrier.

## CONCLUSION

In this study, we have explored various aspects of the nsp16 MTase activation and function. MD simulations were set up to generate conformational and dynamics information of nsp16 in different nsp10 and ligand binding states. Comparisons between different MD setups revealed how the flexible loops of the nsp16 were modulated by the nsp10 to form its SAM pocket in order to carry out the MTase function. AI/ML methods were then applied to identify the meaning molecular states and establish the binding pathway/mechanism of the nsp16/nsp10 complex.

The nsp10 binding is essential for the nsp16 MTase function in SARS-CoV-2. (11, 12, 18) In this study, we identified the mechanism of nsp16 SAM binding pocket reacting to the nsp10 binding. Since the part of SAMBL1 overlapped with the binding interface, the binding of nsp10 significantly altered the backbone fluctuation. In absence of nsp10, the nsp16 SAMBL1 residue, Gly6869, backbone dihedral flipped and caused the following protein backbone now overlapping with the SAM pocket. Notably, the Gly6869 residue is also conserved across pathogenic Coronaviruses. This result explained at the molecular level the activation of nsp16 enzymatic activity upon nsp10 binding.

The RNA binding loops host the viral RNA for the methylation process. (17) Our result revealed that these loops were affected by the ligands in the nsp16 SAM pocket. The GL1 acted as a door that swing open with no binding ligands. The simulation replicas ubiquitously agreed on the open pose after 500 ns with empty SAM pocket. After binding with SAM/SAH/SFG, the GL1 acted in a non-structural fashion. The GL2 was stabilized with SAM as the MTase reactant as a result of a stronger hydrogen bond forming between the ligand ribose and protein backbone.

The SAM pocket itself was intended for SAM binding and SAH exiting. Our simulation also recorded the SAH exiting process and disclosed the selective binding property of protein, differentiated by the ligand property. The SAM pocket was stabilized upon binding with nsp10, providing a more suitable binding environment for more rigid SAM, which was less favoring to flexible SAH. As a result, the nsp16/nsp10 complex form was able to maintain binding with SAM and decoupling with SAH automatically.

In the binding analysis, we identified a key residue, nsp10 Leu4298, which sit in the hydrophobic pocket during the native binding state. During the binding process, it first latched into the same pocket to ensure correct binding pose between the nsp16 and nsp10, which were subsequently formed, following the rearrangement of the interfacial loops. The binding approach undertook by the nsp16/nsp10 complex locked in the binding interface with part of the binding regions on the loops prior to the full contact. The flexibility of these loops allowed these partial binding to happen, and left enough adjustability for the rest of native contacts to form. Therefore, the binding process was guided through the early contact of flexible region at the interface.

In summary, we explored the detailed interactions within nsp16/nsp10 complex with MD simulations and AI/ML analyses. These results explained the mechanism of nsp16 (de)activation and part of its function in a molecular level. It can help to drive understanding of nsp16 MTase machinery in the coronavirus family, and provide new perspectives for designing more effective inhibitors for nsp16 function, that would prompt early immune detection of viral infection. Moreover, this knowledge is also applicable in general with protein function activation and corroborative modulation.

## AUTHOR CONTRIBUTIONS

HM, MRL, KS and AR designed the research. HM carried out the simulations. HM, MRL and AB contributed analysis and visualization tools for the simulation datasets. HM, AB, AR and SCC developed data analytics modules for the simulations. All authors contributed to the writing and editing of this paper.

## ACKNOWLEDGMENTS

This research was supported by the Exascale Computing Project (17-SC20-SC), a collaborative effort of the US DOE Office of Science and the National Nuclear Security Administration, the National Institute of Allergy and Infectious Diseases, National Institutes of Health Award Number P01AI165077 (AR), the National Science Foundation Award Number 2117896 and supported by the DOE through the National Virtual Biotechnology Laboratory, a consortium of DOE national laboratories focused on response to COVID-19, with funding from the Coronavirus CARES Act. KS is grateful for the funding from the Chicago Biomedical Consortium COVID-19 Award 60057467 and from HHS/NIH/NIAID contracts HHSN272201700060C and 75N93022C00035.

## SUPPLEMENTARY MATERIAL

An online supplement to this article can be found by visiting BJ Online at http://www.biophysj.org.

## REFERENCES

1. Dong, E., H. Du, and L. Gardner, 2020. An interactive web-based dashboard to track COVID-19 in real time. The Lancet infectious diseases 20:533–534.

2. CDC: Centers for Disease Control and Prevention, 2024. COVID Data Tracker. https://covid.cdc.gov/covid-data-tracker.

3. Meganck, R. M., and R. S. Baric, 2021. Developing therapeutic approaches for twenty-first-century emerging infectious viral diseases. Nature medicine 27:401–410.

4. Xiu, S., A. Dick, H. Ju, S. Mirzaie, F. Abdi, S. Cocklin, P. Zhan, and X. Liu, 2020. Inhibitors of SARS-CoV-2 entry: current and future opportunities. Journal of medicinal chemistry 63:12256–12274.

5. Snijder, E., E. Decroly, and J. Ziebuhr, 2016. The nonstructural proteins directing coronavirus RNA synthesis and processing. Advances in virus research 96:59–126.

6. da Silva, S. J. R., C. T. Alves da Silva, R. P. G. Mendes, and L. Pena, 2020. Role of nonstructural proteins in the pathogenesis of SARS-CoV-2. Journal of Medical Virology 92:1427–1429.

7. Vicenti, I., M. Zazzi, and F. Saladini, 2021. SARS-CoV-2 RNA-dependent RNA polymerase as a therapeutic target for COVID-19. Expert Opinion on Therapeutic Patents 31:325–337.

8. Daffis, S., K. J. Szretter, J. Schriewer, J. Li, S. Youn, J. Errett, T.-Y. Lin, S. Schneller, R. Zust, H. Dong, et al., 2010. 2-O methylation of the viral mRNA cap evades host restriction by IFIT family members. Nature 468:452–456.

9. Ramanathan, A., G. B. Robb, and S.-H. Chan, 2016. mRNA capping: biological functions and applications. Nucleic acids research 44:7511–7526.

10. Decroly, E., I. Imbert, B. Coutard, M. Bouvet, B. Selisko, K. Alvarez, A. E. Gorbalenya, E. J. Snijder, and B. Canard, 2008. Coronavirus nonstructural protein 16 is a cap-0 binding enzyme possessing (nucleoside-2 O)-methyltransferase activity. Journal of virology 82:8071–8084.

11. Chen, Y., C. Su, M. Ke, X. Jin, L. Xu, Z. Zhang, A. Wu, Y. Sun, Z. Yang, P. Tien, et al., 2011. Biochemical and structural insights into the mechanisms of SARS coronavirus RNA ribose 2-O-methylation by nsp16/nsp10 protein complex. PLoS pathogens 7:e1002294.

12. Chen, Y., and D. Guo, 2016. Molecular mechanisms of coronavirus RNA capping and methylation. Virologica Sinica 31:3–11.

13. Menachery, V. D., K. Debbink, and R. S. Baric, 2014. Coronavirus non-structural protein 16: evasion, attenuation, and possible treatments. Virus research 194:191–199.

14. Schindewolf, C., and V. D. Menachery, 2023. Coronavirus 2-O-methyltransferase: A promising therapeutic target. Virus Research 336:199211. https://www.sciencedirect.com/science/article/pii/S0168170223001739.

15. Bergant, V., S. Yamada, V. Grass, Y. Tsukamoto, T. Lavacca, K. Krey, M. Mühlhofer, S. Wittmann, A. Ensser, A. Herrmann, A. vom Hemdt, Y. Tomita, S. Matsuyama, T. Hirokawa, Y. Huang, A. Piras, C. A. Jakwerth, M. Oelsner, S. Thieme, A. Graf, S. Krebs, H. Blum, B. M. Kümmerer, A. Stukalov, C. B. Schmidt-Weber, M. Igarashi, T. Gramberg, A. Pichlmair, and H. Kato, 2022. Attenuation of SARS-CoV-2 replication and associated inflammation by concomitant targeting of viral and host cap 2’-O-ribose methyltransferases. The EMBO Journal 41:e111608. https://www.embopress.org/doi/abs/10.15252/embj.2022111608.

16. Bouvet, M., A. Lugari, C. C. Posthuma, J. C. Zevenhoven, S. Bernard, S. Betzi, I. Imbert, B. Canard, J.-C. Guillemot, P. Lécine, et al., 2014. Coronavirus Nsp10, a critical co-factor for activation of multiple replicative enzymes. Journal of Biological Chemistry 289:25783–25796.

17. Rosas-Lemus, M., G. Minasov, L. Shuvalova, N. L. Inniss, O. Kiryukhina, J. Brunzelle, and K. J. Satchell, 2020. High-resolution structures of the SARS-CoV-2 2-O-methyltransferase reveal strategies for structure-based inhibitor design. Science signaling 13:eabe1202.

18. Vithani, N., M. D. Ward, M. I. Zimmerman, B. Novak, J. H. Borowsky, S. Singh, and G. R. Bowman, 2021. SARS-CoV-2 Nsp16 activation mechanism and a cryptic pocket with pan-coronavirus antiviral potential. Biophysical Journal 120:2880–2889.

19. Bouvet, M., C. Debarnot, I. Imbert, B. Selisko, E. J. Snijder, B. Canard, and E. Decroly, 2010. In vitro reconstitution of SARS-coronavirus mRNA cap methylation. PLoS pathogens 6:e1000863.

20. Decroly, E., C. Debarnot, F. Ferron, M. Bouvet, B. Coutard, I. Imbert, L. Gluais, N. Papageorgiou, A. Sharff, G. Bricogne, et al., 2011. Crystal structure and functional analysis of the SARS-coronavirus RNA cap 2-O-methyltransferase nsp10/nsp16 complex. PLoS pathogens 7:e1002059.

21. Minasov, G., M. Rosas-Lemus, L. Shuvalova, N. L. Inniss, J. S. Brunzelle, C. M. Daczkowski, P. Hoover, A. D. Mesecar, and K. J. F. Satchell, 2021. Mn2+ coordinates Cap-0-RNA to align substrates for efficient 2′-<i>O</i>-methyl transfer by SARS-CoV-2 nsp16. Science Signaling 14:eabh2071. https://www.science.org/doi/abs/10.1126/scisignal.abh2071.

22. Pan, J., X. Peng, Y. Gao, Z. Li, X. Lu, Y. Chen, M. Ishaq, D. Liu, M. L. DeDiego, L. Enjuanes, et al., 2008. Genome-wide analysis of protein-protein interactions and involvement of viral proteins in SARS-CoV replication. PloS one 3:e3299.

23. Rogstam, A., M. Nyblom, S. Christensen, C. Sele, V. O. Talibov, T. Lindvall, A. A. Rasmussen, I. André, Z. Fisher, W. Knecht, et al., 2020. Crystal structure of non-structural protein 10 from severe acute respiratory syndrome coronavirus-2. International journal of molecular sciences 21:7375.

24. Kubatova, N., N. Qureshi, N. Altincekic, R. Abele, J. Bains, B. Ceylan, J. Ferner, C. Fuks, B. Hargittay, M. Hutchison, et al., 2021. 1H, 13C, and 15N backbone chemical shift assignments of coronavirus-2 non-structural protein Nsp10. Biomolecular NMR Assignments 15:65–71.

25. Ma, Y., L. Wu, N. Shaw, Y. Gao, J. Wang, Y. Sun, Z. Lou, L. Yan, R. Zhang, and Z. Rao, 2015. Structural basis and functional analysis of the SARS coronavirus nsp14–nsp10 complex. Proceedings of the National Academy of Sciences 112:9436–9441.

26. Wang, H., S. R. Rizvi, D. Dong, J. Lou, Q. Wang, W. Sopipong, Y. Su, F. Najar, P. K. Agarwal, F. Kozielski, and S. Haider, 2023. Emerging variants of SARS-CoV-2 NSP10 highlight strong functional conservation of its binding to two non-structural proteins, NSP14 and NSP16. eLife 12:RP87884. 10.7554/eLife.87884.

27. Kremling, V., S. Falke, Y. Fernández-García, C. Ehrt, A. Kiene, B. Klopprogge, E. Scheer, F. Barthels, P. Middendorf, S. Kühn, S. Günther, M. Rarey, H. N. Chapman, D. Oberthür, and J. Sprenger, 2024. SARS-CoV-2 methyltransferase nsp10-16 in complex with natural and drug-like purine analogs for guiding structure-based drug discovery 10.7554/eLife.98310.1.

28. Eastman, P., J. Swails, J. D. Chodera, R. T. McGibbon, Y. Zhao, K. A. Beauchamp, L.-P. Wang, A. C. Simmonett, M. P. Harrigan, C. D. Stern, et al., 2017. OpenMM 7: Rapid development of high performance algorithms for molecular dynamics. PLoS computational biology 13:e1005659.

29. Lindorff-Larsen, K., S. Piana, K. Palmo, P. Maragakis, J. L. Klepeis, R. O. Dror, and D. E. Shaw, 2010. Improved side-chain torsion potentials for the Amber ff99SB protein force field. Proteins: Structure, Function, and Bioinformatics 78:1950–1958.

30. Wang, J., W. Wang, P. A. Kollman, and D. A. Case, 2006. Automatic atom type and bond type perception in molecular mechanical calculations. Journal of molecular graphics and modelling 25:247–260.

31. Wang, J., R. M. Wolf, J. W. Caldwell, P. A. Kollman, and D. A. Case, 2004. Development and testing of a general amber force field. Journal of computational chemistry 25:1157–1174.

32. Darden, T., D. York, and L. Pedersen, 1993. Particle mesh Ewald: An N log (N) method for Ewald sums in large systems. The Journal of chemical physics 98:10089–10092.

33. Case, D. A., H. M. Aktulga, K. Belfon, I. Ben-Shalom, S. R. Brozell, D. S. Cerutti, T. E. Cheatham III, V. W. D. Cruzeiro, T. A. Darden, R. E. Duke, et al., 2021. Amber 2021. University of California, San Francisco.

34. Bhowmik, D., S. Gao, M. T. Young, and A. Ramanathan, 2018. Deep clustering of protein folding simulations. BMC bioinformatics 19:47–58.

35. Abadi, M., A. Agarwal, P. Barham, E. Brevdo, Z. Chen, C. Citro, G. S. Corrado, A. Davis, J. Dean, M. Devin, S. Ghemawat, I. Goodfellow, A. Harp, G. Irving, M. Isard, Y. Jia, R. Jozefowicz, L. Kaiser, M. Kudlur, J. Levenberg, D. Mané, R. Monga, S. Moore, D. Murray, C. Olah, M. Schuster, J. Shlens, B. Steiner, I. Sutskever, K. Talwar, P. Tucker, V. Vanhoucke, V. Vasudevan, F. Viégas, O. Vinyals, P. Warden, M. Wattenberg, M. Wicke, Y. Yu, and X. Zheng, 2015. TensorFlow: Large-Scale Machine Learning on Heterogeneous Systems. https://www.tensorflow.org/, software available from tensorflow.org.

36. Pedregosa, F., G. Varoquaux, A. Gramfort, V. Michel, B. Thirion, O. Grisel, M. Blondel, P. Prettenhofer, R. Weiss, V. Dubourg, J. Vanderplas, A. Passos, D. Cournapeau, M. Brucher, M. Perrot, and E. Duchesnay, 2011. Scikit-learn: Machine Learning in Python. Journal of Machine Learning Research 12:2825–2830.

37. Sculley, D., 2010. Web-scale k-means clustering. In Proceedings of the 19th international conference on World wide web. 1177–1178.

38. Blondel, V. D., J.-L. Guillaume, R. Lambiotte, and E. Lefebvre, 2008. Fast unfolding of communities in large networks. Journal of statistical mechanics: theory and experiment 2008:P10008.

39. Newman, M. E., 2006. Modularity and community structure in networks. Proceedings of the national academy of sciences 103:8577–8582.

40. Gerstein, M., and C. Chothia, 1996. Packing at the protein-water interface. Proceedings of the National Academy of Sciences 93:10167–10172.

41. Walker, R. C., M. F. Crowley, and D. A. Case, 2008. The implementation of a fast and accurate QM/MM potential method in Amber. Journal of computational chemistry 29:1019–1031.

