## Supporting Information for "Mechanistic insights into the association and activation of the SARS-CoV-2 2’-O-Methyltransferase (NSP16)"

Manuscript submitted to **Biophysical** *Journal*

**Article**

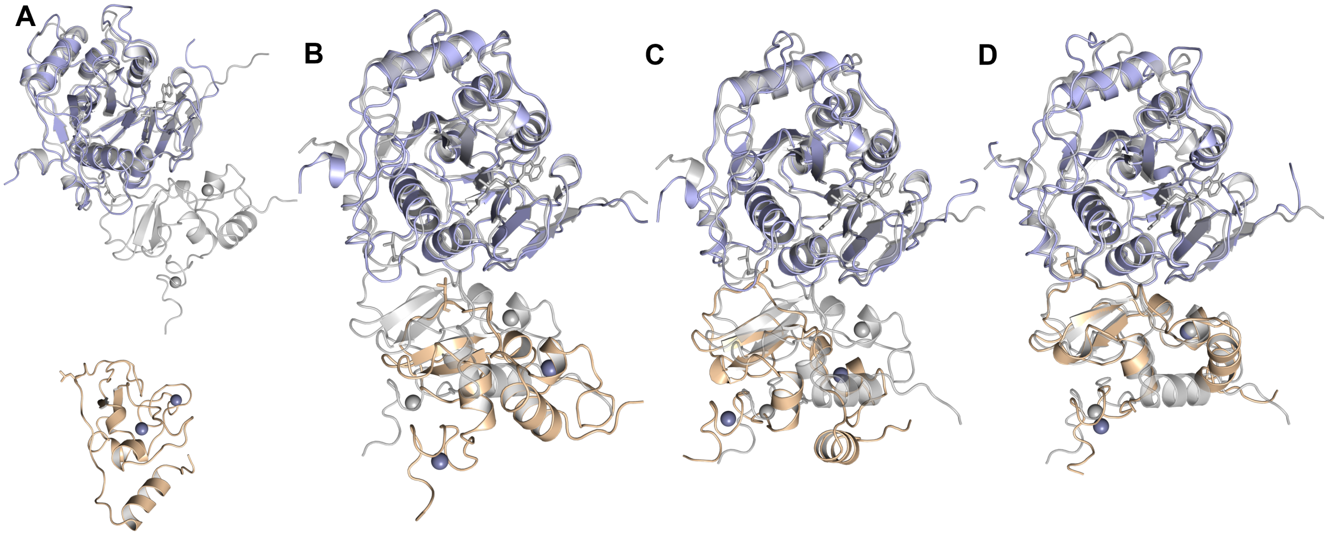


Figure S1: The conformational states along the binding pathway, the separated, transition 2, transition 1 and binding states from left to right. The proteins initiated binding process via nsp10 Leu4298 docked in the nsp16 hydrophobic pocket, to ensure the native binding interface. The process was then completed with the formation of the H-bonds between Leu4298 and nsp16 Gln6885.

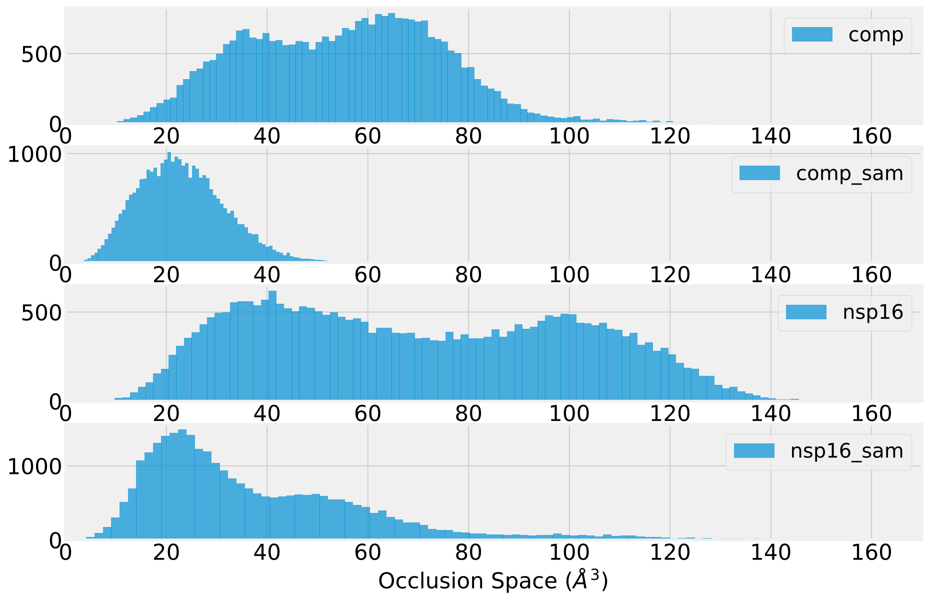


Figure S2: The histogram of occlusion space from simulation trajectories from apo-nsp16/nsp10 complex, nsp16-SAM/nsp10 complex, apo-nsp16 and nsp16-SAM monomer..

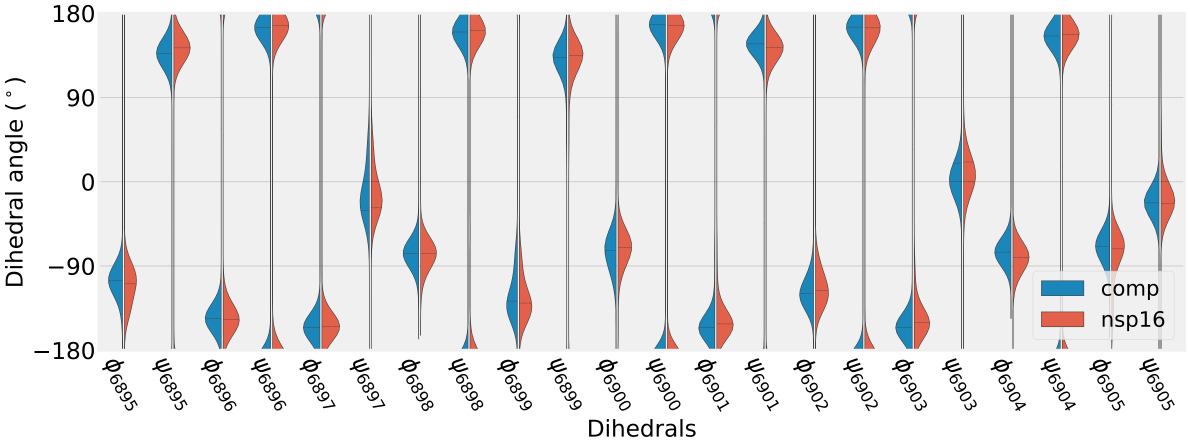


Figure S3: The histogram of nsp16 SAMBL2 backbone dihedral angles from nsp16/nsp10 complex and nsp16 monomer. All were in the same configuration between the complex and monomer form.

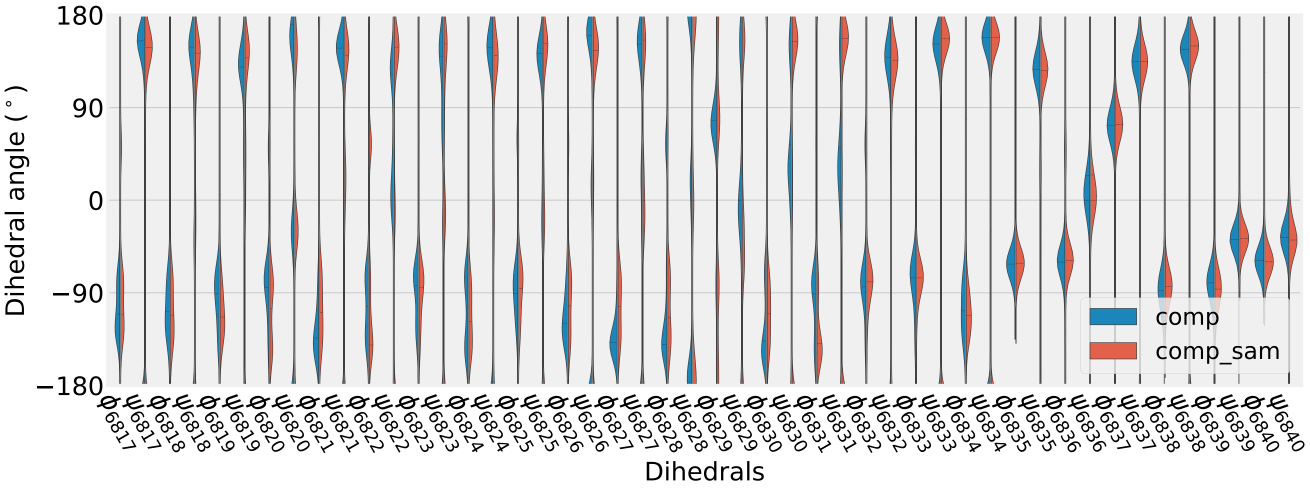


Figure S4: The histogram of nsp16 GL1 backbone dihedral angles from nsp16/nsp10 with and without SAM molecule.

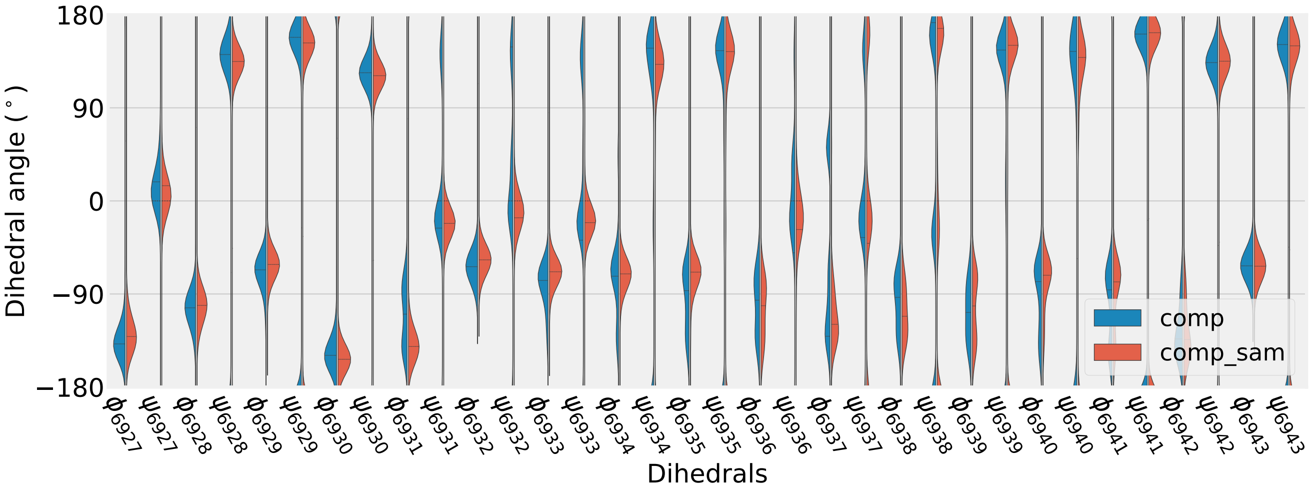


Figure S5: The histogram of nsp16 GL2 backbone dihedral angles from nsp16/nsp10 with and without SAM molecule.

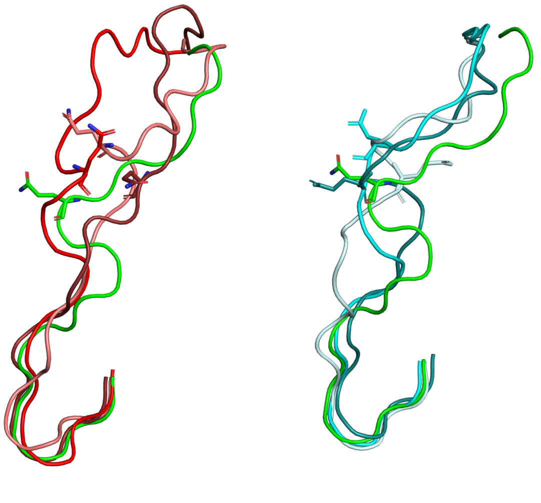

Figure S6: The GL1 configuration of nsp16/nsp10 complex binding with SAH (left) and SFG (right).

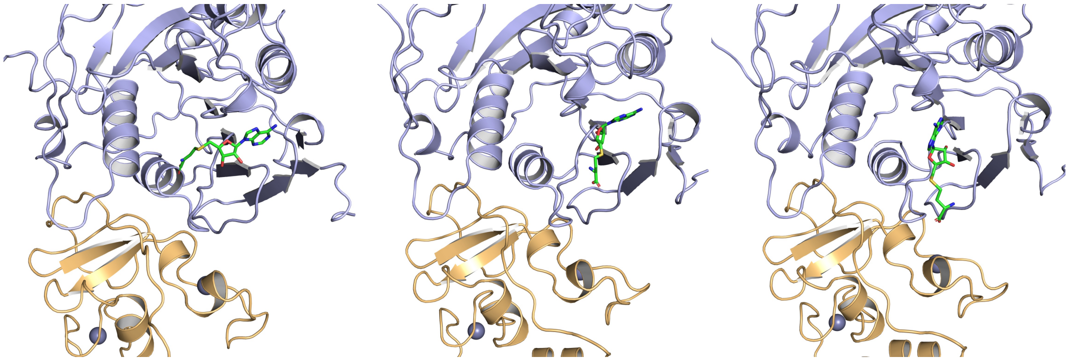

Figure S7: The escape of SAH from the SAM pocket of nsp16/nsp10 complex.

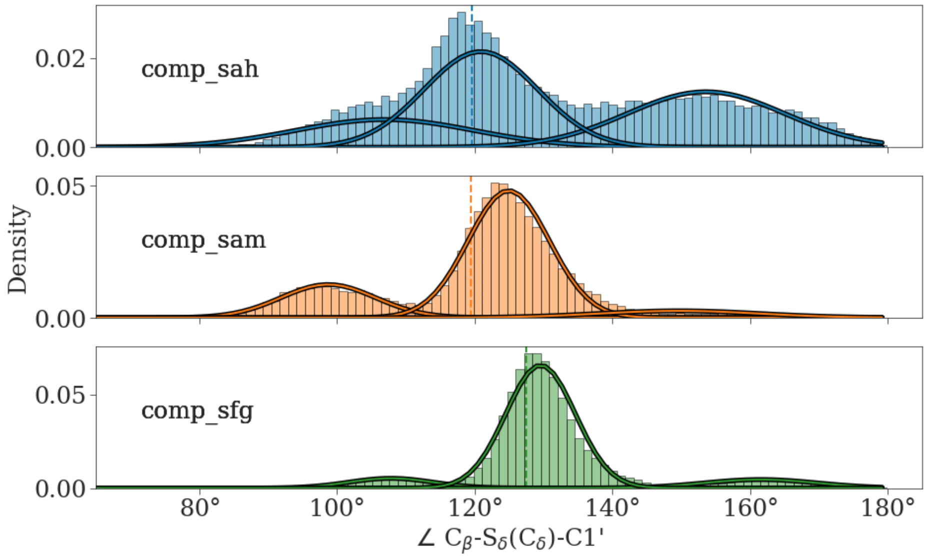


Figure S8: The histogram of the ligand angle, $\angle C_\beta - S_\eta(C_\eta) - C'$, in the SAM pocket of nsp16/nsp10 complex form during MD simulations.
